# A NOVEL PREGNANT RAT MODEL FOR LABOR INDUCTION AND AUGMENTATION WITH OXYTOCIN

**DOI:** 10.1101/2021.08.11.455293

**Authors:** Tusar Giri, Jia Jiang, Zhiqiang Xu, Ronald Mccarthy, Carmen M. Halabi, Eric Tycksen, Alison G. Cahill, Sarah K. England, Arvind Palanisamy

## Abstract

**Background:** Despite the widespread use of oxytocin for induction of labor, mechanistic insights into maternal and neonatal wellbeing are lacking because of the absence of an animal model that recapitulates modern obstetric practice.

**Objective:** The objectives of this research were to create and validate a hi-fidelity animal model that mirrors labor induction with oxytocin in parturients and to assess its translational utility.

**Study Design:** The study was performed in timed-pregnant Sprague Dawley dams. The model consisted of a subcutaneously implanted microprocessor-controlled infusion pump on gestational day 18 that was pre-programmed to deliver an escalating dose of intravenous oxytocin on gestational day 21 to induce birth. Once predictable delivery of healthy pups was achieved, we validated the model with molecular biological experiments on the uterine myometrium and telemetry-supported assessment of changes in intrauterine pressure. Finally, we applied this model to test the hypothesis that labor induction with oxytocin was associated with oxidative stress in the newborn brain with a comprehensive array of biomarker assays and oxidative stress gene expression studies.

**Results:** During the iterative model development phase, we confirmed the optimal gestational age for pump implantation, the concentration of oxytocin, and the rate of oxytocin administration. Exposure to anesthesia and surgery during pump implantation was not associated with significant changes in the cortical transcriptome. Activation of pump with oxytocin on gestational day 21 resulted in predictable delivery of pups within 8-12 hours. Increased frequency of change of oxytocin infusion rate was associated with dystocic labor. Labor induction and augmentation with oxytocin was associated with increased expression of the oxytocin receptor gene in the uterine myometrium, decreased expression of the oxytocin receptor protein on the myometrial cell membrane, and cyclical increases in intrauterine pressure. Examination of the frontal cortex of vaginally delivered newborn pups born after oxytocin-induced labor did not reveal an increase in oxidative stress compared to saline-treated control pups. Specifically, there were no significant changes in oxidative stress biomarkers involving both the oxidative stress (reactive oxygen/nitrogen species, 4-hydroxynonenal, protein carbonyl) and the antioxidant response (total glutathione, total antioxidant capacity). In addition, there were no significant differences in the expression of 16 genes emblematic of the oxidative stress response pathway.

**Conclusions:** Collectively, we provide a viable and realistic animal model for labor induction and augmentation with oxytocin. We demonstrate its utility in addressing clinically relevant questions in obstetric practice that could not be mechanistically ascertained otherwise. Based on our findings, labor induction with oxytocin is not likely to cause oxidative stress in the fetal brain. Adoption of our model by other researchers would enable new lines of investigation related to the impact of perinatal oxytocin exposure on the mother-infant dyad.

## INTRODUCTION

Labor induction and augmentation with oxytocin (Oxt) is one of the most prevalent clinical interventions in modern obstetric practice(1–5). Despite widespread use for over 50 years, most research has focused on the contractile effects of Oxt and associated obstetric outcomes(4–7). Whether Oxt affects the fetus remains sparsely studied, despite controversial epidemiological evidence suggesting a link between the use of Oxt and neurodevelopmental disorders (8–14). Importantly, most preclinical studies that examine this question do so without inducing birth(15–18), making them contextually less germane. An important scientific roadblock is the absence of an animal model that mirrors induction of labor in pregnant women with Oxt, presumably due to the technical difficulty in delivering an incrementally higher dose of intravenous Oxt over time in a free-moving animal. In this report, we surmounted these challenges to create and validate a hi-fidelity pregnant rat model for elective labor induction and augmentation with Oxt using an implantable, programmable, microprocessor-controlled precision drug delivery pump.

## MATERIALS AND METHODS

### Study design

All experiments reported here were approved by the Institutional Animal Care and Use Committee at Washington University in St. Louis (#20170010) and comply with the ARRIVE guidelines. A schematic of the study design is presented in **Fig. 1**.

**Fig. 1.**
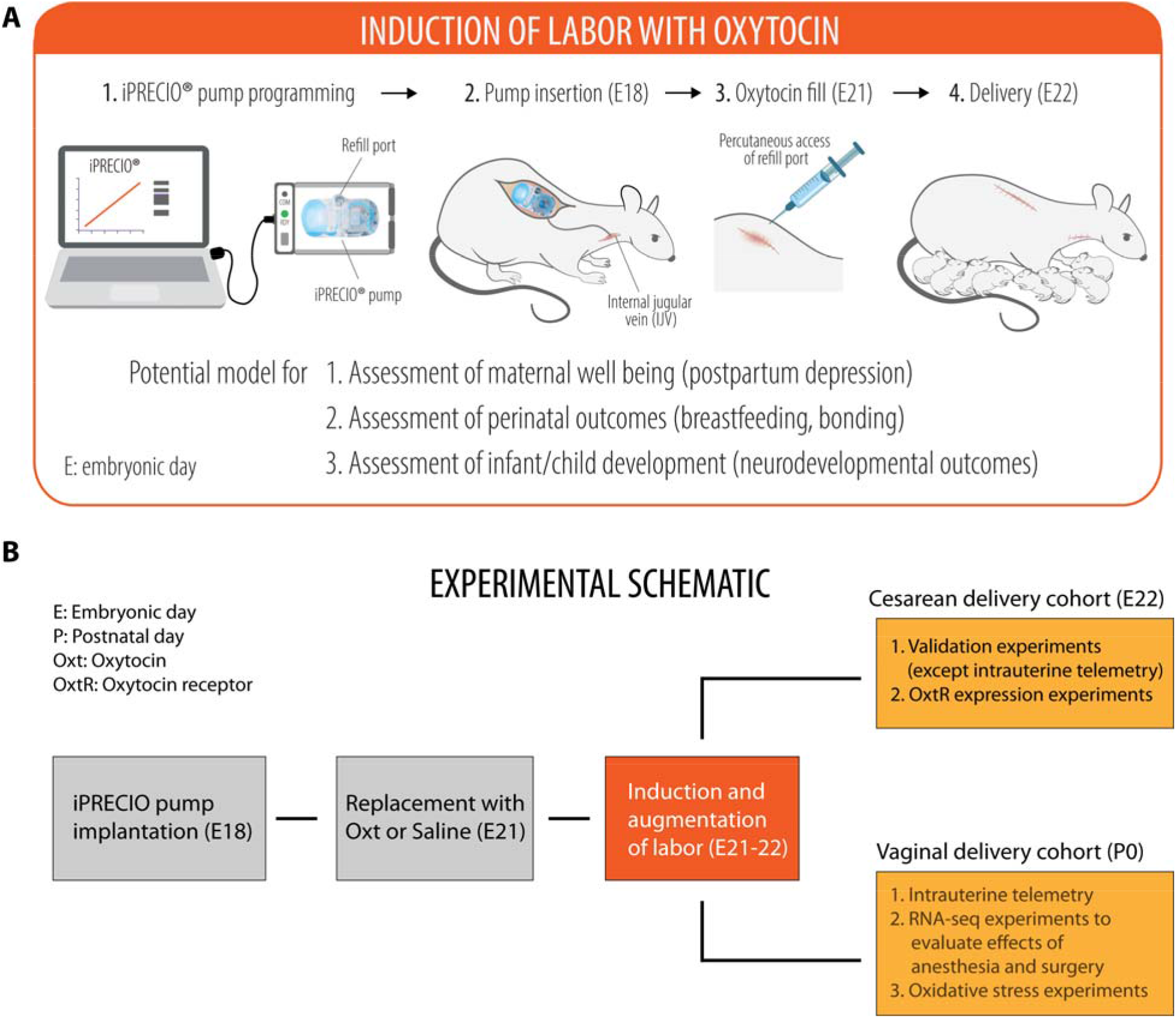
Experimental schematic for labor induction with oxytocin in term pregnant rat. (**A**) A cartoon depicting the programming and implantation of iPRECIO^®^ pump in a pregnant rat followed by birth of healthy pups. (**B**) Experimental schematic showing the overall experimental outline with two separate cohorts for cesarean and vaginal delivery. In the vaginal delivery cohort, there were three sets of independent experiments for (i) intrauterine telemetry, (ii) RNA-seq experiments to assess the impact of in utero exposure to anesthesia and surgery on the cortical transcriptome of the newborn brain, and (iii) examination of oxidative stress in newborn pups after either Oxt or saline, respectively.

### Development of the pregnant rat model for labor induction and augmentation with Oxt

The system consists of a subcutaneously placed iPRECIO^®^ infrared-controlled microinfusion pump (SMP-200, Primetech Corporation) connected to the right internal jugular vein in an embryonic day (E)18 Sprague Dawley dam (Charles River Laboratories) (presented as a photo montage in **Fig. 2**). Briefly, the dam was anesthetized with 2% isoflurane followed by subcutaneous implantation of the iPRECIO^®^ pump approximately 2-3 cm below the nape of the neck and creation of a tunnel to deliver the pump tubing next to the internal jugular vein, into which it was secured in place with ligatures. The reservoir of the iPRECIO^®^ pump was primed with sterile normal saline prior to implantation and was pre-programmed to deliver an infusion rate of 10 μl/h for 72 h to keep the tubing patent until E21. Two hours before completion of the saline infusion at 72 h, the reservoir was accessed subcutaneously under brief isoflurane anesthesia to aspirate the saline and was refilled with 900 μl of Oxt (Selleck Chemicals, 50 μg/mL in normal saline). This was followed by the pre-programmed infusion rate of 5 μl/h for 4 h, 10 μl/h for 4 h, 20 μl/h for 4 h, and 30 μl/h for 12 h (iPRECIO^®^ Management System) (**Fig. 3**).

**Fig. 2.**
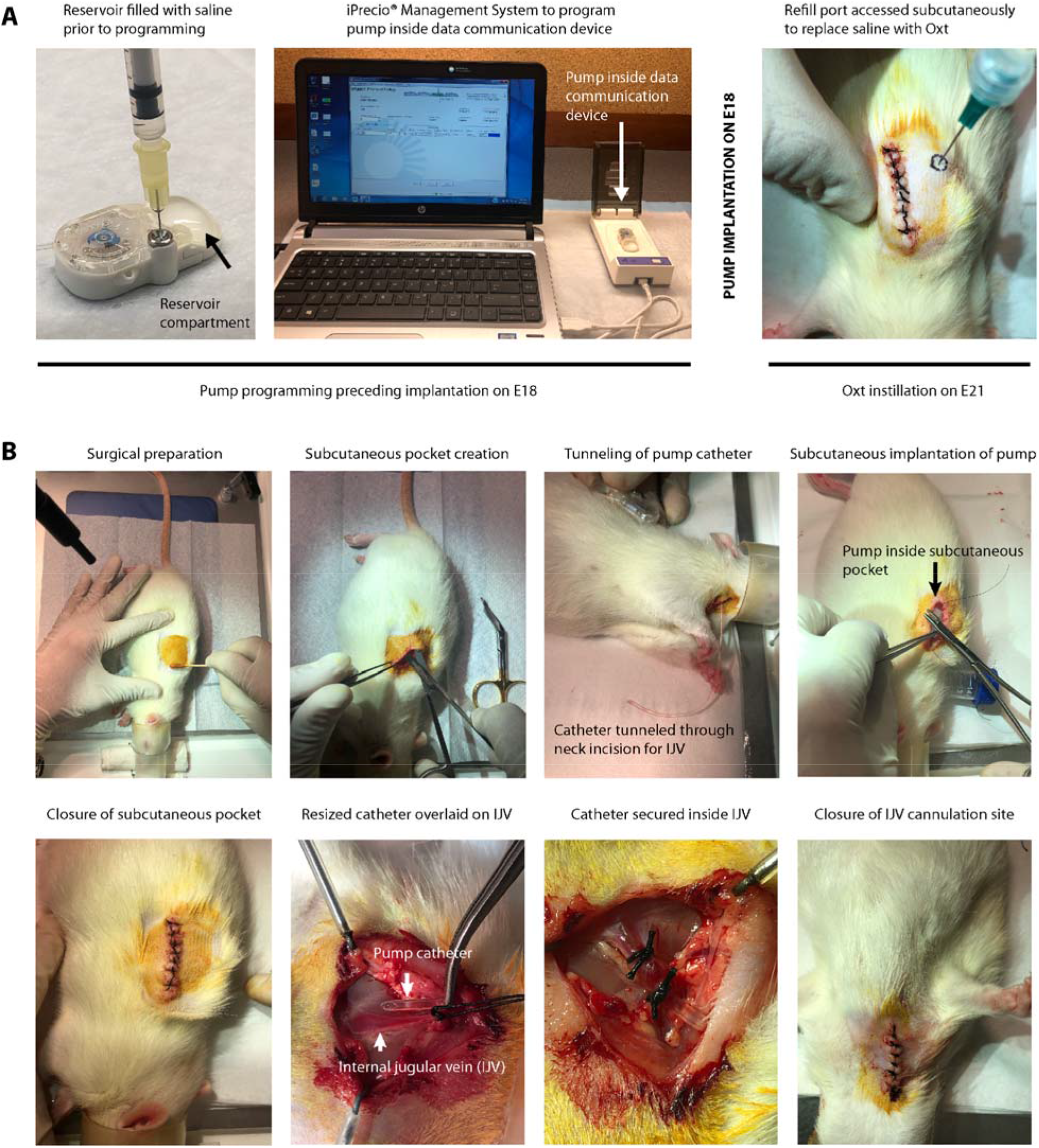
Workflow of experimental and surgical procedures associated with creation of the model. (**A**) Workflow for iPRECIO^®^ pump programming prior to implantation on E18 and subsequent replacement of saline with Oxt by subcutaneous access of the refill port on E21. (**B**) Left to right: sequential surgical workflow for implantation of the pre-programmed iPRECIO^®^ pump followed by internal jugular vein cannulation with the pump catheter on E18. All surgical procedures were performed in strict accordance with institutional guidelines for rodent surgery, anesthesia, and analgesia.

**Fig. 3.**
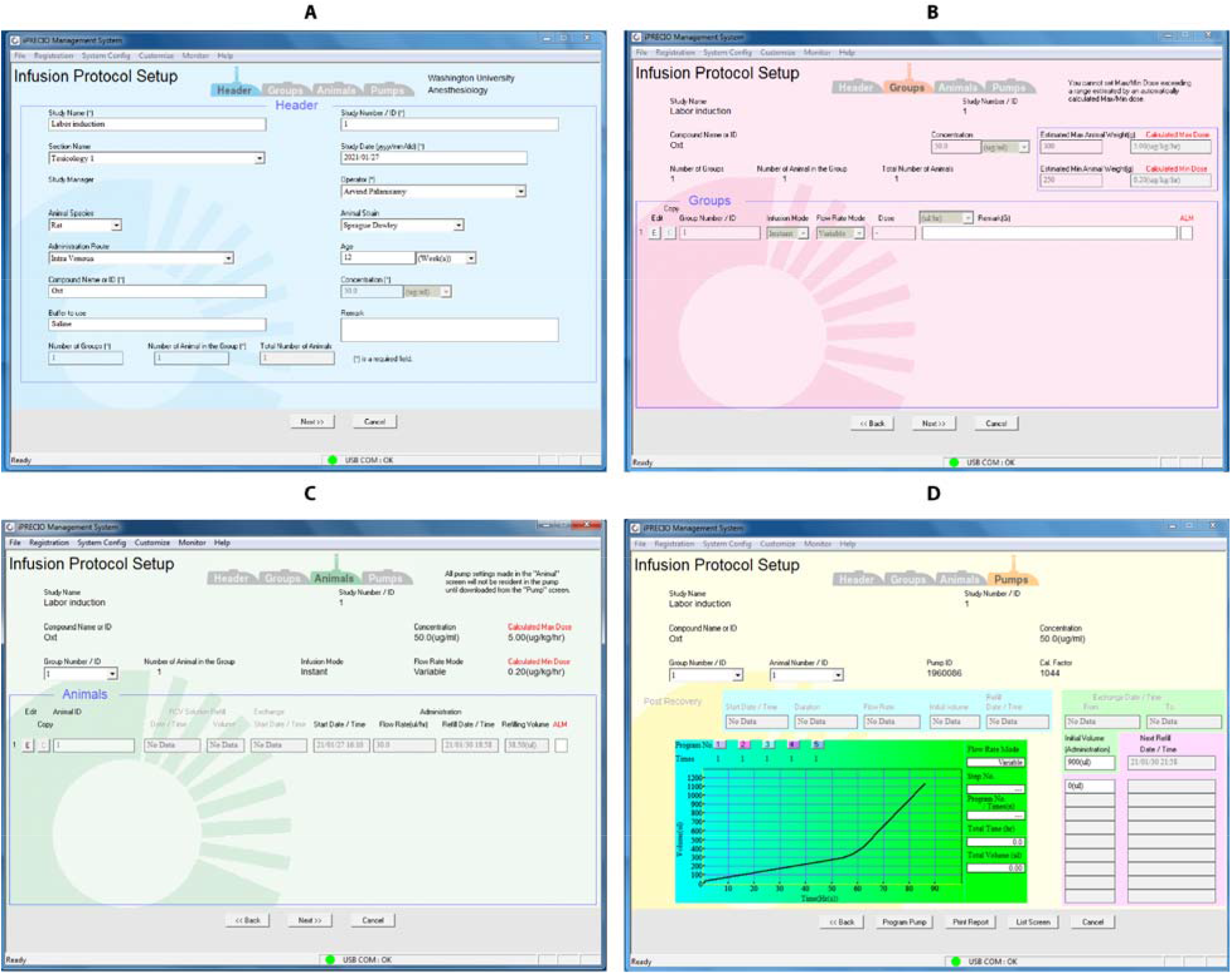
A walkthrough of the sequential steps (A-D) for pump programming with iPRECIO^®^ Management System software. All procedures were performed in accordance with the manufacturer’s instructions. (**A**) Infusion protocol set up page that allows input of all the variables necessary for programming of the pump. (**B-C)** Variable flow rate mode is chosen to ensure gradual escalation of Oxt dose over time. (**D**) A graphic representation of Oxt dosing showing the volume that is to be administered over time. The pump ID and calibration factor is automatically detected and inputted by the data communication device.

### Validation experiments

Though the witnessed birth of pups offered functional validation, we examined the effect of Oxt on the uterine myometrium with molecular biological assays, immunohistochemistry, and telemetric assessment of changes in uterine pressure.

#### (i) OxtR gene expression

Briefly, approximately 0.5 cm x 1 cm rectangular piece of myometrial tissue was harvested from the anti-mesometrial aspect of the uterus after 8-12 h of exposure to either Oxt (100 mcg/mL concentration) or saline. Sample processing and OxtR qPCR was performed with a custom TaqMan^®^ OxtR probe as described by us previously(19).

#### (ii) Western blot for OxtR expression

Membrane-associated proteins were isolated from approximately 100 mg of uterine myometrial tissue using Mem-PER Plus Membrane Protein Extraction Kit (catalog# 89842, ThermoFisher Scientific, Inc.) following manufacturer’s instructions and subjected to immunoblotting with appropriate positive and negative controls (Cat#: LY400333, Origene Technologies, Inc). Details are provided in the Supplementary Materials and Methods.

#### (iii) Immunohistochemistry

Briefly, 5-μm frozen sections of uterine myometrium embedded in OCT compound were obtained using Leica CM1510 S cryostat and immunostained for phosphorylated myosin light chain kinase (1:200 rabbit anti-mouse phosphomyosin light chain kinase, Invitrogen) and imaged with the Zeiss Axioskop 40 microscope. OxtR protein expression was assessed by immunostaining with goat anti-rat OXTR antibody (1:100; Origene) and revealed with Alexa Fluor^®^ 594 labeled rabbit anti-goat antibody (1:300, Invitrogen). All primary antibodies were incubated overnight at 4°C followed by a 1 h incubation with secondary antibodies at room temperature. Imaging was performed with Olympus BX60 fluorescence microscope with designated filter sets.

#### (iv) Uterine telemetry

To assess whether initiation of Oxt was temporally associated with increase in intrauterine pressure, we performed pressure recordings with telemetry as described previously by us for mice (20, 21). Briefly, under isoflurane anesthesia and sterile precautions, we inserted a pressure catheter in the right horn between the uterine wall and the fetus under sterile precautions during pump implantation in E18 dams. To minimize the possibility that telemetry recordings could represent spontaneous labor, we advanced the time of replacement of saline with Oxt to 48 h instead of 72 h (i.e., E20 - two days before term gestation). The pressure catheter was connected to a PhysioTel PA-C10 transmitter (Data Sciences International) placed in the lower portion of the abdominal cavity. Telemetry recordings were performed at 500 Hz with Dataquest ART data acquisition system version 4.10 (DSI) sampling every 5 min for 15 sec intervals for 6 h at baseline, followed by recordings 48 h later when Oxt was initiated and continued until the birth of the pups.

### Effect of *in utero* exposure to anesthesia and surgery on the neonatal cortical transcriptome

To rule out the possibility of adverse effects on the fetal brain from intrauterine exposure to anesthesia and surgery during pump implantation(22, 23), we examined the cortical transcriptome of newborn pups delivered spontaneously by unhandled *vs*. surgically implanted dams. Briefly, 2 brains from spontaneously delivered newborn pups of either sex were collected within 2 h of birth from 6 dams (n=3 each for spontaneous labor and saline-filled iPRECIO^®^ pump at E18). Total RNA was extracted from the right cerebral cortex using RNAeasy kit (Qiagen) and subjected to RNA-seq (Genome Technology Access Center core facility). Only RNA with RIN > 9.5 were used for RNA-seq. Processing of samples, sequencing, and analysis were done as described by us previously (19) and in the Supplementary Materials and Methods.

### Assessment of biomarkers of oxidative stress in the newborn brain

Oxt-induced cyclical uterine contractions cause lipid peroxidative injury(24), decrease the anti-oxidant glutathione in cord blood(25), and increase amniotic fluid lactate(26, 27) suggesting the possibility of oxidative stress. Because the developing fetal/neonatal brain is vulnerable to oxidative stress (28), we used our model to investigate this question. Briefly, brains were isolated from vaginally delivered newborn pups immediately after birth, snap frozen, and stored at −80°C for oxidative stress assays. Cortical lysates were prepared according to the assay type and protein concentration was determined using BCA Protein Assay Kit (ThermoFisher Scientific) prior to the assays. All assays were performed in duplicate, and fluorescence/absorbance was read with Tecan Infinite^®^ M200 PRO multimode plate reader using appropriate filter sets as recommended by the manufacturer. We assayed for total free radicals (OxiSelect™ In Vitro ROS/RNS Assay Kit, #STA-347), 4-hydroxynonenal (lipid peroxidation marker, OxiSelect™ HNE Adduct Competitive ELISA Kit, # STA-838), protein carbonyl (marker of oxidative damage to proteins; OxiSelect™ Protein Carbonyl ELISA kit, # STA-310), total glutathione (OxiSelect™ Total Glutathione Assay kit, # STA-312), and total antioxidant capacity (OxiSelect™ TAC Assay Kit, # STA-360). All assays were purchased from Cell BioLabs, Inc (San Diego, CA).

### Expression of genes mediating oxidative stress in the newborn brain

From the same set of experiments as above, brains were isolated from additional pups born after exposure to either Oxt or saline (n= 6-8 per group), snap frozen, and immediately stored at - 80°C. Processing of total RNA for gene expression experiments was performed as described by us previously(19). Expression levels of 16 genes relevant to oxidative stress (*Mtnd2*, *Mtnd5*, *Mtcyb*, *Mt-co1*, *Mt-atp8*) and antioxidant (*Sod1*, *Sod2*, *Gpx1*, *Gpx4*, *Prdx1*, *Cat*, *Gsr*, *Nox3*, *Nox4*, *Txnip*, *Txrnd2*) pathways were assayed in duplicate along with four endogenous housekeeping control genes (*18S rRNA*, *Gapdh*, *Pgk1*, and *Actb*) and reported as described previously(19).

### Statistical analysis

Data outliers were detected and eliminated using ROUT (robust regression and outlier analysis) with Q set to 10%. Because our pilot experiments with a higher dose of Oxt (100 mcg/mL concentration) showed no sex differences in the expression of oxidative stress markers in the newborn brain, all subsequent analyses were performed regardless of sex of the offspring. RNA-seq data were analyzed as described by us previously(19). Quantitative data were analyzed with Welch’s t-test with p ≤ 0.05 considered significant, while oxidative stress gene expression data were analyzed with unpaired student’s t-test followed by Bonferroni correction with an adjusted p-value ≤ 0.003 considered significant. All analyses, with the exception of RNA-seq data, were performed on Prism 8 for Mac OS X (Graphpad Software, Inc, La Jolla, CA) and expressed as mean ± S.E.M.

## RESULTS

### Development of the model for labor induction with Oxt

Overall, 44 timed-pregnant Sprague Dawley dams were used for the study (Supplementary Table S1). A video walkthrough of the experimental setup is presented as Supplementary Movie S1. With the final regimen for Oxt as described in Methods, dams gave birth to pups predictably within 8-12 h. Litter size and weight gain trajectory of the offspring from one experimental cohort are presented in Supplementary Table S2. Handling of critical steps and troubleshooting are described in greater detail in the Supplementary Materials and Methods.

### Validation of the model

The best validation of our model was the successful vaginal delivery of thriving pups within 12 h after initiation of the Oxt regimen (Supplementary Movie S2). In addition, we confirmed the presence of immunoreactive phosphorylated myosin light chain kinase (MLCK)(29), a serine/threonine kinase and a downstream regulator of the effects of Oxt on the actin-myosin ATPase, in Oxt-exposed myometrium (**Fig. 4A**). Next, we confirmed that Oxt initiation was accompanied by a rise in intrauterine pressure, a *sine qua non* feature of labor(30–32), and lasting until birth of all pups (**Fig. 4B**). This was associated with an increase in OxtR gene expression in the uterine myometrium (**Fig. 4C**). In contrast, exposure to Oxt for at least 8 h resulted in a decrease in OxtR immunoreactivity (**Fig. 4D**) and membrane bound OxtR protein expression (**Fig. 4E**) similar to human data. Collectively, we established the translational relevance of our model by mirroring both Oxt management of labor and its effect on the uterine myometrium.

**Fig. 4.**
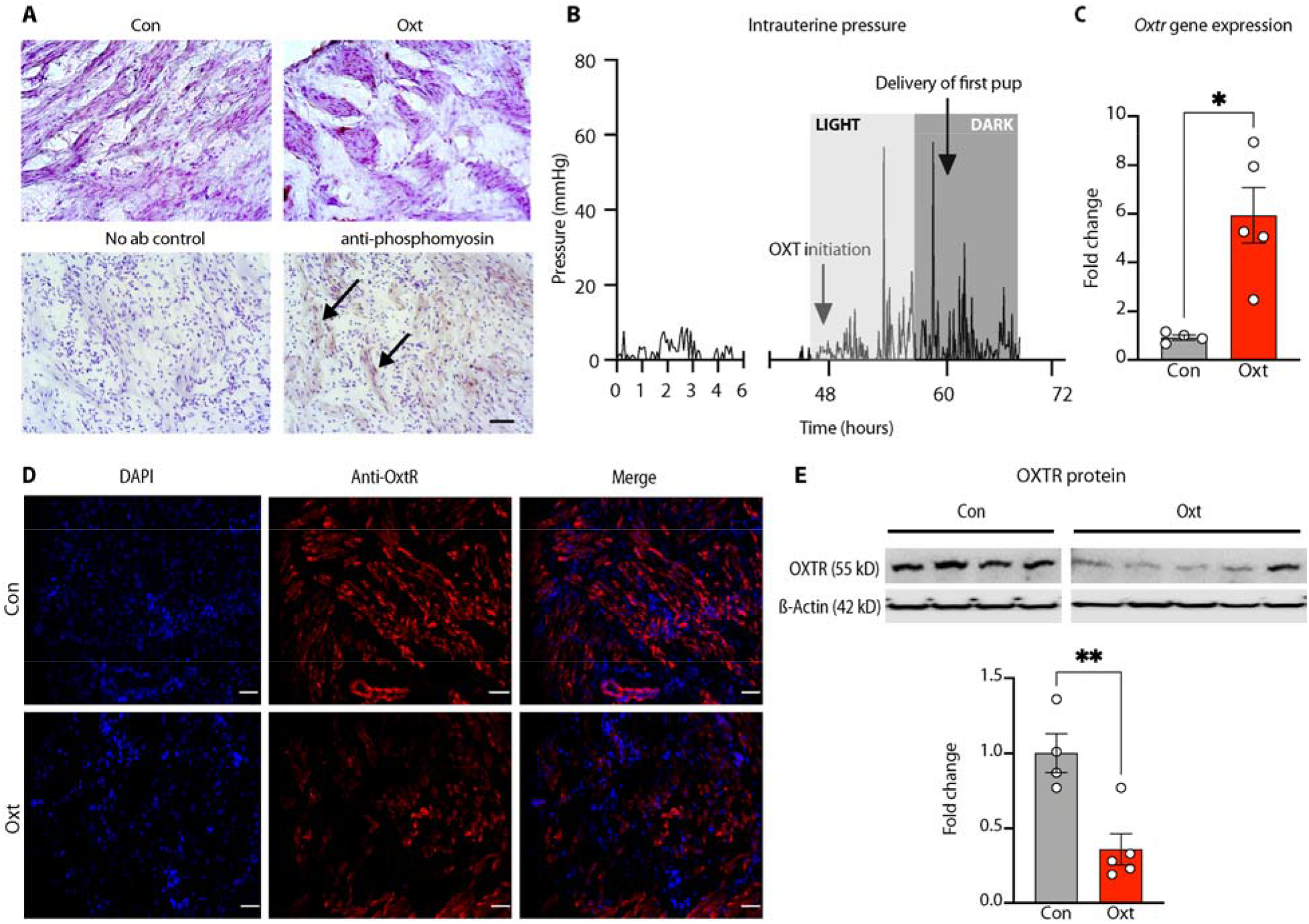
Validation of the model for labor induction with oxytocin. (**A**) Visualization of uterine contraction. Upper panel: uterine myometrium harvested from pregnant E21 rats at least 8 h after either saline (control) or intravenous Oxt infusion and stained with hematoxylin. 20x photomicrographs showing lack of clustering of uterine myocytes in saline-treated myometrium (left) compared to extensive clustering in the Oxt-exposed myometrium (right). Lower panel: 5-μm frozen sections from Oxt-exposed myometrium stained without (left) or with rabbit anti-mouse phosphomyosin light chain kinase showing prominent staining among clustered uterine myocytes revealed with anti-rabbit HRP conjugate (marked by arrows) (right). Nuclei counterstained with hematoxylin. Scale bar = 100μM. (**B**) Labor induction with Oxt causes cyclical increases in intrauterine pressure. Labor was induced with Oxt at 48 h after pump implantation (around 12 noon, light cycle, E20) and intrauterine pressure changes were monitored with telemetry. Oxt initiation was associated with acute and cyclical increases in intrauterine pressure until birth (first pup delivered around 21:00 h, dark cycle, E20). Light and dark cycle from 07:00-19:00 and 19:00-07:00, respectively. (**C**) Labor induction with Oxt was associated with a significant increase in OxtR gene expression at 8-12 h. (**D**) Labor induction with Oxt decreases OxtR immunoreactivity in the rat uterus. Sample 20x photomicrographs from 5-µm sections of the uterine myometrium stained with goat anti-rat OxtR antibody (1:100) and revealed with Alexa Fluor^®^ 594 labeled rabbit anti-goat antibody (1:300). Note naïve uterine myometrium with bright staining for OxtR in the upper panel, in sharp contrast to Oxt-exposed myometrial tissue in the lower panel where staining was scant, suggesting downregulation of OxtR. Scale bar = 50μM. (**E**) Representative western blot showing a decrease in membrane associated OxtR protein expression after labor induction with Oxt and quantified with densitometry. Data analyzed with Welch’s t-test and presented as mean ± SEM; *p ≤ 0.05, **p ≤ 0.01.

### Effect of surgery and anesthesia during pump implantation on the developing brain

Because exposure to anesthesia and surgery can affect the developing brain(22, 23, 33–35), we compared the cortical transcriptomes of newborn pups born to spontaneously laboring dams that were not exposed to pump implantation surgery *vs*. those that were implanted with a saline-filled pump on E18 (and therefore requiring anesthesia). Unbiased RNA-seq analyses of the cerebral cortex of vaginally delivered newborn pups revealed no significant changes in the cortical transcriptome after exposure to surgery and anesthesia as shown by the lack of significantly differentially expressed genes in the volcano plot (**Fig. 5A**; heat map in Supplementary Fig. S1.). Principal component analysis (**Fig. 5B**) revealed that the major source of variance was not the treatment condition but the sex of the offspring, albeit not significant. Top up- and downregulated genes from GO and KEGG analyses are presented in **Fig. 5C–E**. A comprehensive list of differentially expressed genes and unadjusted p-value significant differentially expressed genes is provided in Supplementary Data S1 and S2, respectively.

**Fig. 5.**
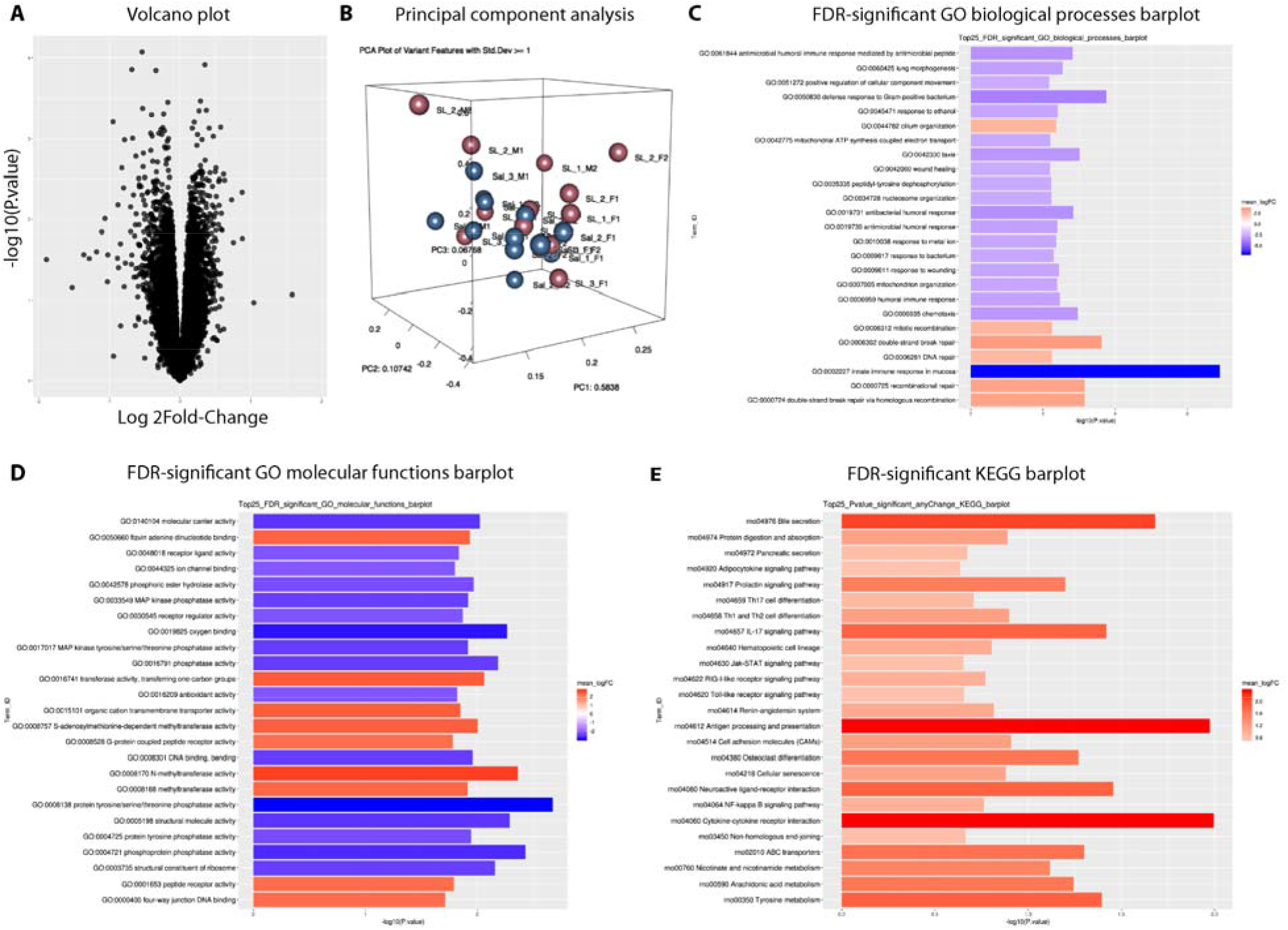
Impact of anesthesia and surgery on the newborn cortical transcriptome. (**A**) Volcano plot showing the absence of significantly differentially expressed genes between the spontaneous labor vs. saline pump groups (n = 2 pups of each sex/dam from 3 dams/treatment condition). (**B**) Principal component analysis (PCA) showing that the major source of variance is not the treatment condition (fuchsia: spontaneous labor, blue: saline pump) but the sex of the offspring, albeit not significant. (**C**-**D**) Top 25 false discovery rate-adjusted significantly up- and downregulated genes for Gene Ontology (GO) biological processes (C) and molecular functions (D) after labor induction with Oxt. (**E**) Significantly upregulated genes with Kyoto Encyclopedia of Genes and Genomes (KEGG) analysis. GO and KEGG analyses revealed a differential impact of anesthesia and surgery on multiple pathways, mostly related to oxygen binding and the immune response, respectively. Therefore, for the rest of our experiments, we used saline pump-implanted dams that eventually labored spontaneously as controls, instead of dams that labored spontaneously without exposure to anesthesia and surgery.

### Examination of the redox state of the fetal cortex after labor induction with Oxt

Labor induction with Oxt was not associated with changes in the concentration of total free radicals, 4-hydroxynonenal or protein carbonyl, in the newborn cortex. Nor were there any significant differences in antioxidant capacity; both glutathione and total antioxidant capacity were unchanged after Oxt (**Fig. 6A**). Furthermore, we did not observe any significant changes in the expression of emblematic genes pertinent to the oxidative stress/antioxidant pathway (**Fig. 6B;** TaqMan qPCR probe list in Supplementary Table S3). Collectively, these data provide reassurance that the use of Oxt for labor induction is unlikely to be associated with oxidative stress in the fetal brain.

**Fig. 6.**
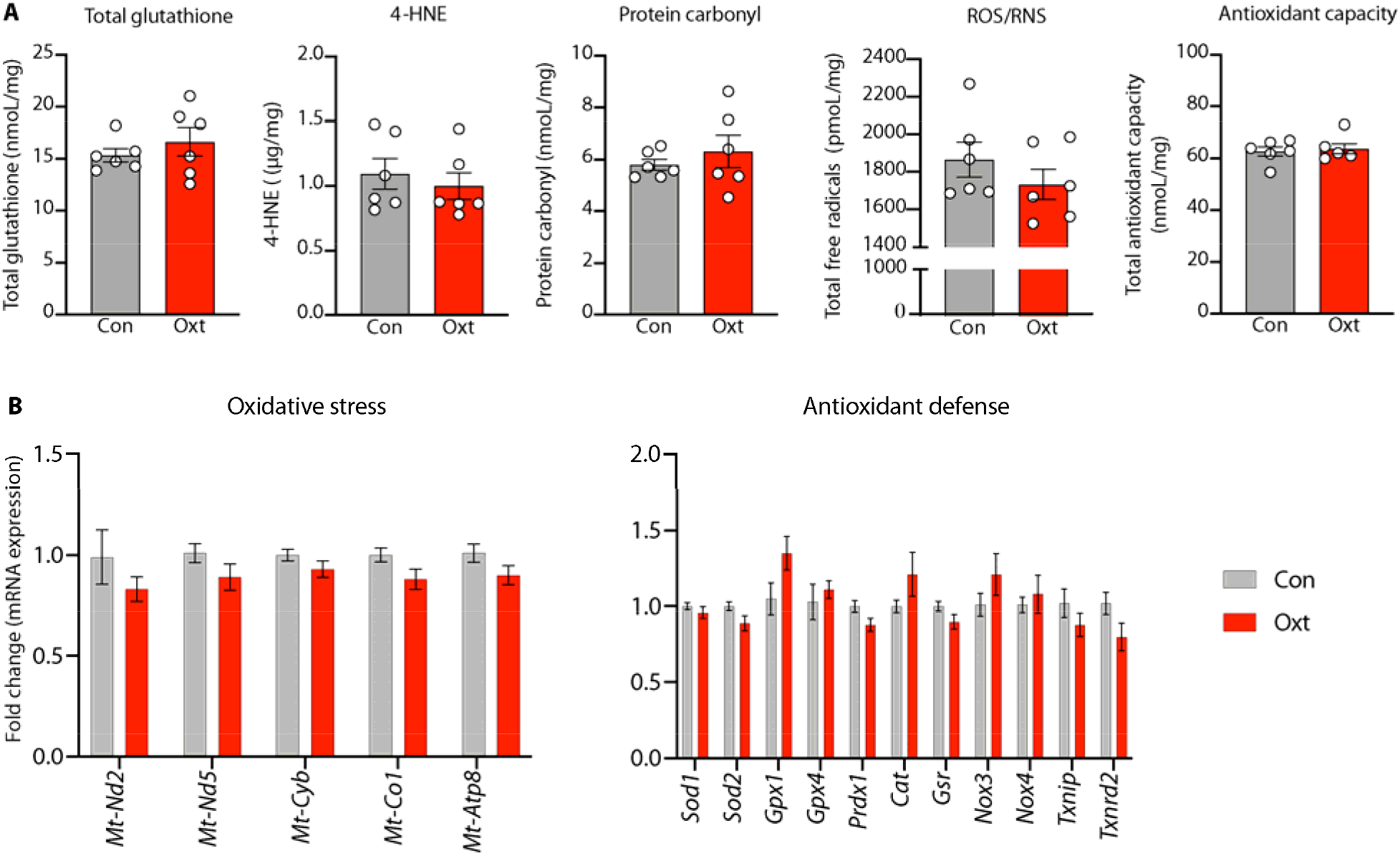
Labor induction with oxytocin is not associated with oxidative stress in the developing brain. (**A**) From left to right: Labor induction with oxytocin was not associated with an increase in either glutathione, 4-hydroxynonenal, protein carbonyl, reactive oxygen/nitrogen species (ROS/RNS), or total antioxidant capacity in the developing fetal cortex. All data are presented per mg of brain protein. (**B**) Expression of genes mediating oxidative stress or antioxidant defense were not significantly differentially expressed in the fetal cortex after labor induction with oxytocin. Collectively, these data indicate that labor induction with Oxt is unlikely to be associated with oxidative stress in the developing brain. Data were analyzed with Welch’s t-test and expressed as mean ± SEM (n=5-8 per treatment condition with pups represented from all unique dams).

## COMMENT

### Principal Findings

Here, we present a realistic and tractable animal model for labor induction with Oxt. In addition to functional validation of the model, we were able to demonstrate features consistent with the use of Oxt in human labor: (i) a decrease in OxtR protein expression in the uterine myometrium, and (ii) confirmation of increased intrauterine pressure with Oxt. Furthermore, we provide evidence for the translational utility of the model by showing that labor induction with Oxt was not associated with oxidative stress in the fetal brain.

### Results in the Context of What is Known

Regarding use of Oxt to induce birth, the only other relevant preclinical study is that of Hirayama et al. which used an osmotic pump to deliver a continuous subcutaneous infusion of Oxt in pregnant mice(36). However, the experimental paradigm did not allow for escalation of Oxt dose nor assessment of the impact of Oxt administration on the uterine myometrium. Another study examined the impact of intravenous Oxt infusion on the fetal brain response to hypoxia/anoxia and showed that pre-conditioning with Oxt increased the concentration of lactate in the fetal brain but reduced the level of malondialdehyde, a lipid peroxidation marker(17). Nevertheless, this study was designed to study the effect of Oxt on the brain adaptation to hypoxia and not to assess the impact of Oxt on the process of birthing. Furthermore, Oxt was administered as a constant infusion, unlike the gradually escalating rate used in our study. These differences perhaps explain why we did not observe a decrease in oxidative stress in the fetal brain. As a G protein-coupled receptor that is sensitive to downregulation, our findings of reduced membrane bound OxtR protein expression after Oxt exposure is broadly consistent with published data in human studies(6). However, to our surprise, expression of the OxtR gene was significantly increased after labor induction with Oxt. We believe that these apparently contradictory findings could be due to the choice of myometrial samples by Phaneuf et al.(6); samples were collected from patients who underwent cesarean delivery after dystocic labor with Oxt suggesting the possibility of abnormal transcription during intrapartum arrest of labor. In contrast, we performed cesarean delivery during uncomplicated labor to facilitate sample collection. This line of thought is supported by the 4-5-fold increased myometrial expression of OxtR gene during uncomplicated labor in rodents(37, 38).

### Clinical Implications

Our findings were reassuring in that even after 8-12 h of exposure to Oxt-induced uterine contractions, there was no evidence for oxidative stress in the newborn brain. Lack of oxidative stress after prolonged exposure to repetitive Oxt-induced uterine contractions in a species in which labor typically lasts between 90-120 min(39), gives us more confidence that this is unlikely to be a concern for the human fetus. Because of the wide variability in Oxt use across the world(40), future research should focus on altering the dose regimens to determine if some of the clinical observations related to oxidative stress are due to differences in Oxt dosing.

### Research Implications

Ethical and logistic challenges significantly limit the scope of mechanistic research on pregnant women and their newborn. Our contextually relevant animal model, by providing unrivaled access to maternal and fetal tissue, has wide-ranging implications for translational research related to perinatal Oxt exposure. This makes our model well suited to investigate lingering concerns about the impact of Oxt on neurobehavioral development of the offspring(8, 14, 41, 42), epigenetic regulation of OxtR in the fetal brain(18), relationship between intrapartum Oxt use and breastfeeding success(43–45), and the complex association between Oxt and postpartum depression(46–48). Furthermore, by scaling down with appropriate equipment (iPRECIO^®^ SMP 310-R with a dedicated wireless communication device), transgenic mouse models could be used to investigate complex gene-environment interaction studies in the perinatal period. Ongoing studies in our laboratory are focused on the transfer of maternally administered Oxt across the placental and fetal blood-brain barriers, and its impact on Oxt-ergic signaling in the fetal brain. Because of the critical importance of Oxt-ergic signaling for satiety and appetite regulation(49), we are also particularly interested in the impact of perinatal Oxt exposure on childhood obesity.

### Strengths and Limitations

The biggest strength of our model is how it mirrors labor induction with Oxt in clinical practice. We prefer not to anthropomorphize our study because biological validation of the effect of oxytocin with the birth of living pups was our motivation. However, the cumulative Oxt dose until birth of the pups, approximately 3-7 μg, is comparable to the dose ranges typically used during human labor. For example, parturients receive on average, a cumulative Oxt dose of 2000-4000 mIU or 2-4 IU (IU = International Unit) during the course of labor (50). Because 1 IU = 1.68 μg of Oxt peptide, this would translate to approximately 3.4-6.8 μg of Oxt, similar to what we used in our model. Considering that our model simulates clinical practice to a large extent, research knowledge generated using this model is more likely to provide reliable and actionable mechanistic data than other currently available models.

Our research has a few limitations. First, our model can be perceived as contrived. Considering the technical challenges of delivering an escalating dose of intravenous Oxt in a free-moving animal to simulate obstetric practice, we considered all possibilities before pursuing this model. Importantly, our model is in no way more traumatic or less realistic than the unilateral carotid artery ligation/ anoxia model to investigate perinatal asphyxia in rodents (51, 52). Second, even though our low-dose Oxt infusion for the first 4 h would have resulted in cervical ripening as demonstrated in laboring women (53), we are unable to provide objective evidence to support that assumption. Nevertheless, because birth of the pups occurred predictably, it is likely a moot concern. Third, we did not compare the extent to which Oxt increases intrauterine pressure compared to saline. Because we had biological validation of pup birth, our objective was to capture the temporal relationship between the initiation of Oxt and the rise in intrauterine pressure rather than assess differences in intrauterine pressure between Oxt-induced and spontaneous labor that have been characterized previously(54).

## Conclusions

In conclusion, we provide a viable and realistic animal model for labor induction and augmentation with Oxt and demonstrate its utility in addressing clinically relevant questions in obstetric practice. Adoption of our model by other researchers would enable new lines of investigation related to the impact of perinatal Oxt exposure on the mother-infant dyad.

## Acknowledgments

None.

## Author contributions

Arvind Palanisamy: Conceptualization, Investigation, Methodology, Project Administration, Resources, Software, Data Curation, Formal analysis, Supervision, Visualization, Writing
Tusar Giri: Investigation (model creation, molecular biology experiments), Methodology, Validation, Writing
Jia Jiang: Investigation (model creation)
Zhiqiang Xu: Investigation (immunohistochemistry experiments)
Ron McCarthy: Investigation (intrauterine telemeter placement)
Carmen M. Halabi: Funding acquisition, Resources (telemetry monitoring), Writing
Sarah K. England: Funding acquisition, Methodology, Resources, Writing
Eric Tycksen: Data curation, Software, Formal analysis (RNA-seq data), Visualization, Writing
Alison G. Cahill. Writing – original draft, review & editing.

## Data sharing statement

The equipment needed to establish the model are commercially available and non-proprietary. All data needed to evaluate the conclusions in the paper are present in the paper and/or the Supplementary Materials. The RNA-seq data used in this publication have been deposited in NCBI’s Gene Expression Omnibus (GEO) and are accessible through the GEO Series accession number GSE161122 (https://www.ncbi.nlm.nih.gov/geo/query/acc.cgi?acc=GSE161122).

## Supplementary Files

### 1. Supplementary Materials and Methods

### 2. Supplementary Figures

**Fig. S1.** RNA-seq data showing the heatmap of differentially expressed genes after *in utero* exposure to anesthesia and surgery for pump implantation.

### 3. Supplementary Tables

**Table S1.** Animal use data.

**Table S2.** Litter data and weight gain trajectory of the offspring.

**Table S3.** Taqman qPCR probe list from ThermoFisher Scientific, Inc.

### 4. Supplementary Movies

**Movie S1.** Experimental set up for iPRECIO^®^ pump implantation. A video walk-through of the overall surgical set up for performing the experiments.

**Movie S2.** Appropriate and nurturing care of the newborn after low dose (50 mcg/mL) Oxt regimen.

**Movie S3.** Poor maternal self-care and pup neglect in a dam implanted with iPRECIO^®^ pump at E20 and treated with high dose of Oxt (100 mcg/mL).

### 5. Supplementary Data Files

**Data S1.** RNA-seq data showing the list of significantly expressed genes in the developing cortex after *in utero* exposure to anesthesia and surgery.

**Data S2.** RNA-seq data showing the list of false discovery rate-unadjusted significantly expressed genes in the developing cortex after *in utero* exposure to anesthesia and surgery.

## References

1. Obstetrics ACoPB--. ACOG Practice Bulletin No. 107: Induction of labor. Obstet Gynecol. 2009;114(2 Pt 1):386–97.

2. Osterman MJ, Martin JA. Recent declines in induction of labor by gestational age. NCHS Data Brief. 2014(155):1–8.

3. Martin JA, Hamilton BE, Osterman MJK, Driscoll AK, Drake P. Births: Final Data for 2016. Natl Vital Stat Rep. 2018;67(1):1–55.

4. Grobman WA, Rice MM, Reddy UM, Tita ATN, Silver RM, Mallett G, et al. Labor Induction versus Expectant Management in Low-Risk Nulliparous Women. N Engl J Med. 2018;379(6):513–23.

5. Souter V, Painter I, Sitcov K, Caughey AB. Maternal and newborn outcomes with elective induction of labor at term. Am J Obstet Gynecol. 2019;220(3):273 e1–e11.

6. Phaneuf S, Rodriguez Linares B, TambyRaja RL, MacKenzie IZ, Lopez Bernal A. Loss of myometrial oxytocin receptors during oxytocin-induced and oxytocin-augmented labour. J Reprod Fertil. 2000;120(1):91–7.

7. Crane JM, Young DC, Butt KD, Bennett KA, Hutchens D. Excessive uterine activity accompanying induced labor. Obstet Gynecol. 2001;97(6):926–31.

8. Gregory SG, Anthopolos R, Osgood CE, Grotegut CA, Miranda ML. Association of autism with induced or augmented childbirth in North Carolina Birth Record (1990-1998) and Education Research (1997-2007) databases. JAMA Pediatr. 2013;167(10):959–66.

9. Weisman O, Agerbo E, Carter CS, Harris JC, Uldbjerg N, Henriksen TB, et al. Oxytocin-augmented labor and risk for autism in males. Behav Brain Res. 2015;284:207–12.

10. Friedlander E, Feldstein O, Mankuta D, Yaari M, Harel-Gadassi A, Ebstein RP, et al. Social impairments among children perinatally exposed to oxytocin or oxytocin receptor antagonist. Early Hum Dev. 2017;106-107:13–8.

11. Kurth L, Haussmann R. Perinatal Pitocin as an early ADHD biomarker: neurodevelopmental risk? J Atten Disord. 2011;15(5):423–31.

12. Guastella AJ, Cooper MN, White CRH, White MK, Pennell CE, Whitehouse AJO. Does perinatal exposure to exogenous oxytocin influence child behavioural problems and autistic-like behaviours to 20 years of age? J Child Psychol Psychiatry. 2018;59(12):1323–32.

13. Oberg AS, D’Onofrio BM, Rickert ME, Hernandez-Diaz S, Ecker JL, Almqvist C, et al. Association of Labor Induction With Offspring Risk of Autism Spectrum Disorders. JAMA Pediatr. 2016;170(9):e160965.

14. Soltys SM, Scherbel JR, Kurian JR, Diebold T, Wilson T, Hedden L, et al. An association of intrapartum synthetic oxytocin dosing and the odds of developing autism. Autism. 2020;24(6):1400–10.

15. Boer GJ. Chronic oxytocin treatment during late gestation and lactation impairs development of rat offspring. Neurotoxicol Teratol. 1993;15(6):383–9.

16. Boer GJ, Kruisbrink J. Effects of continuous administration of oxytocin by an accurel device on parturition in the rat and on development and diuresis in the offspring. J Endocrinol. 1984;101(2):121–9.

17. Boksa P, Zhang Y, Nouel D. Maternal Oxytocin Administration Before Birth Influences the Effects of Birth Anoxia on the Neonatal Rat Brain. Neurochem Res. 2015;40(8):1631–43.

18. Kenkel WM, Perkeybile AM, Yee JR, Pournajafi-Nazarloo H, Lillard TS, Ferguson EF, et al. Behavioral and epigenetic consequences of oxytocin treatment at birth. Sci Adv. 2019;5(5):eaav2244.

19. Palanisamy A, Giri T, Jiang J, Bice A, Quirk JD, Conyers SB, et al. In utero exposure to transient ischemia-hypoxemia promotes long-term neurodevelopmental abnormalities in male rat offspring. JCI Insight. 2020;5(10).

20. Pierce SL, Kutschke W, Cabeza R, England SK. In vivo measurement of intrauterine pressure by telemetry: a new approach for studying parturition in mouse models. Physiol Genomics. 2010;42(2):310–6.

21. Rada CC, Pierce SL, Grotegut CA, England SK. Intrauterine telemetry to measure mouse contractile pressure in vivo. Journal of visualized experiments : JoVE. 2015(98):e52541.

22. Palanisamy A. Maternal anesthesia and fetal neurodevelopment. Int J Obstet Anesth. 2012;21(2):152–62.

23. Palanisamy A, Baxter MG, Keel PK, Xie Z, Crosby G, Culley DJ. Rats exposed to isoflurane in utero during early gestation are behaviorally abnormal as adults. Anesthesiology. 2011;114(3):521–8.

24. Calderon TC, Wu W, Rawson RA, Sakala EP, Sowers LC, Boskovic DS, et al. Effect of mode of birth on purine and malondialdehyde in umbilical arterial plasma in normal term newborns. J Perinatol. 2008;28(7):475–81.

25. Schneid-Kofman N, Silberstein T, Saphier O, Shai I, Tavor D, Burg A. Labor augmentation with oxytocin decreases glutathione level. Obstet Gynecol Int. 2009;2009:807659.

26. Murphy M, Butler M, Coughlan B, Brennan D, O’Herlihy C, Robson M. Elevated amniotic fluid lactate predicts labor disorders and cesarean delivery in nulliparous women at term. Am J Obstet Gynecol. 2015;213(5):673 e1–8.

27. Wiberg-Itzel E, Pembe AB, Wray S, Wihlback AC, Darj E, Hoesli I, et al. Level of lactate in amniotic fluid and its relation to the use of oxytocin and adverse neonatal outcome. Acta Obstet Gynecol Scand. 2014;93(1):80–5.

28. Perrone S, Tataranno LM, Stazzoni G, Ramenghi L, Buonocore G. Brain susceptibility to oxidative stress in the perinatal period. J Matern Fetal Neonatal Med. 2015;28 Suppl 1:2291–5.

29. Moore F, Bernal AL. Myosin light chain kinase and the onset of labour in humans. Exp Physiol. 2001;86(2):313–8.

30. Seitchik J, Chatkoff ML. Intrauterine pressure wave form characteristics in hypocontractile labor before and after oxytocin administration. Am J Obstet Gynecol. 1975;123(4):426–34.

31. Seitchik J, Chatkoff ML. Oxytocin-induced uterine hypercontractility pressure wave forms. Obstetrics and gynecology. 1976;48(4):436–41.

32. Bakker PC, Van Rijswijk S, van Geijn HP. Uterine activity monitoring during labor. J Perinat Med. 2007;35(6):468–77.

33. Adhikari A. Distributed circuits underlying anxiety. Front Behav Neurosci. 2014;8:112.

34. Creeley CE, Dikranian KT, Dissen GA, Back SA, Olney JW, Brambrink AM. Isoflurane-induced apoptosis of neurons and oligodendrocytes in the fetal rhesus macaque brain. Anesthesiology. 2014;120(3):626–38.

35. Vutskits L, Xie Z. Lasting impact of general anaesthesia on the brain: mechanisms and relevance. Nat Rev Neurosci. 2016;17(11):705–17.

36. Hirayama T, Hiraoka Y, Kitamura E, Miyazaki S, Horie K, Fukuda T, et al. Oxytocin induced labor causes region and sex-specific transient oligodendrocyte cell death in neonatal mouse brain. J Obstet Gynaecol Res. 2020;46(1):66–78.

37. Helguera G, Eghbali M, Sforza D, Minosyan TY, Toro L, Stefani E. Changes in global gene expression in rat myometrium in transition from late pregnancy to parturition. Physiol Genomics. 2009;36(2):89–97.

38. Shchuka VM, Abatti LE, Hou H, Khader N, Dorogin A, Wilson MD, et al. The pregnant myometrium is epigenetically activated at contractility-driving gene loci prior to the onset of labor in mice. PLoS Biol. 2020;18(7):e3000710.

39. Catheline G, Touquet B, Besson JM, Lombard MC. Parturition in the rat: a physiological pain model. Anesthesiology. 2006;104(6):1257–65.

40. Daly D, Minnie KCS, Blignaut A, Blix E, Vika Nilsen AB, Dencker A, et al. How much synthetic oxytocin is infused during labour? A review and analysis of regimens used in 12 countries. PLoS One. 2020;15(7):e0227941.

41. Gottlieb MM. A Mathematical Model Relating Pitocin Use during Labor with Offspring Autism Development in terms of Oxytocin Receptor Desensitization in the Fetal Brain. Comput Math Methods Med. 2019;2019:8276715.

42. Wahl RU. Could oxytocin administration during labor contribute to autism and related behavioral disorders?--A look at the literature. Med Hypotheses. 2004;63(3):456–60.

43. Jonas W, Johansson LM, Nissen E, Ejdeback M, Ransjo-Arvidson AB, Uvnas-Moberg K. Effects of intrapartum oxytocin administration and epidural analgesia on the concentration of plasma oxytocin and prolactin, in response to suckling during the second day postpartum. Breastfeed Med. 2009;4(2):71–82.

44. Lara-Cinisomo S, McKenney K, Di Florio A, Meltzer-Brody S. Associations Between Postpartum Depression, Breastfeeding, and Oxytocin Levels in Latina Mothers. Breastfeed Med. 2017;12(7):436–42.

45. Uvnas Moberg K, Ekstrom-Bergstrom A, Buckley S, Massarotti C, Pajalic Z, Luegmair K, et al. Maternal plasma levels of oxytocin during breastfeeding-A systematic review. PLoS One. 2020;15(8):e0235806.

46. Jobst A, Krause D, Maiwald C, Hartl K, Myint AM, Kastner R, et al. Oxytocin course over pregnancy and postpartum period and the association with postpartum depressive symptoms. Arch Womens Ment Health. 2016;19(4):571–9.

47. Kroll-Desrosiers AR, Nephew BC, Babb JA, Guilarte-Walker Y, Moore Simas TA, Deligiannidis KM. Association of peripartum synthetic oxytocin administration and depressive and anxiety disorders within the first postpartum year. Depress Anxiety. 2017;34(2):137–46.

48. Thul TA, Corwin EJ, Carlson NS, Brennan PA, Young LJ. Oxytocin and postpartum depression: A systematic review. Psychoneuroendocrinology. 2020;120:104793.

49. Lawson EA. The effects of oxytocin on eating behaviour and metabolism in humans. Nat Rev Endocrinol. 2017;13(12):700–9.

50. Roloff K, Peng S, Sanchez-Ramos L, Valenzuela GJ. Cumulative oxytocin dose during induction of labor according to maternal body mass index. Int J Gynaecol Obstet. 2015;131(1):54–8.

51. Vannucci RC, Connor JR, Mauger DT, Palmer C, Smith MB, Towfighi J, et al. Rat model of perinatal hypoxic-ischemic brain damage. J Neurosci Res. 1999;55(2):158–63.

52. Rumajogee P, Bregman T, Miller SP, Yager JY, Fehlings MG. Rodent Hypoxia-Ischemia Models for Cerebral Palsy Research: A Systematic Review. Front Neurol. 2016;7:57.

53. Ferguson JE, 2nd, Head BH, Frank FH, Frank ML, Singer JS, Stefos T, et al. Misoprostol versus low-dose oxytocin for cervical ripening: a prospective, randomized, double-masked trial. Am J Obstet Gynecol. 2002;187(2):273–9; discussion 9–80.

54. Seitchik J, Chatkoff ML, Hayashi RH. Intrauterine pressure waveform characteristics of spontaneous and oxytocin- or prostaglandin F2alpha--induced active labor. Am J Obstet Gynecol. 1977;127(3):223–7.

