## Supplementary material for "A NOVEL PREGNANT RAT MODEL FOR LABOR INDUCTION AND AUGMENTATION WITH OXYTOCIN": IOL_Model_Supplementary File_Aug10.docx

**1. Supplementary Materials and Methods**

**Study design**

Because of the iterative process involved in the creation of the animal model, we did not predetermine the sample size. However, after confirming the success of the Oxt regimen, sample sizes for all the hypothesis-testing experiments were guided by our previous work on the effect of Oxt-induced uterine hypercontractility on oxidative stress in the fetal brain (Palanisamy et al., JCI Insight 2020). To minimize litter effects, we used one randomly chosen pup per dam when possible.

**Handling of critical steps and troubleshooting**

Ensuring appropriate anesthesia, analgesia, and body temperature, with meticulous attention to sterility and surgical technique was critical for the success of the experiments. The duration of surgery was approximately 30-45 min, and prior to closure of the subcutaneous pocket with 4.0 silk sutures, buprenorphine SR 1mg/kg was injected subcutaneously for post-operative analgesia. Nevertheless, we observed complications related either to the surgery, or the choice of gestational age or the Oxt regimen. One issue we encountered was surgical wound infection despite a single dose of subcutaneous antibiotic (enrofloxacin 10 mg/kg) during the time of pump implantation. Therefore, we changed our antibiotic regimen to extended oral administration via drinking water (trimethoprim /sulfamethoxazole 240 mg/5 mL suspension in 250 mL drinking water). Similarly, suture dehiscence was observed in 3 dams, which we eliminated in subsequent experiments by coating the suture lines with cyanoacrylate glue (Vetbond Tissue Adhesive, 3M). Implantation of the pump at E20 (i.e., 2 days before expected delivery) was associated with a high incidence of either preterm or dystocic labor. Significantly, we also noticed a high rate of either cannibalization, a type of maternal behavior that is known to be uncommon in this species, or poor maternal care with pup neglect (Supplementary Movie S3). Therefore, to minimize this possibility, we advanced the pump implantation day to E18. All pumps were filled with saline, and in the dams assigned to receive Oxt, saline was replaced with Oxt by accessing the reservoir subcutaneously at 72 h . We performed an iterative process for assessing various Oxt concentrations (from 1 mg/mL to 100 mcg/mL to 50 mcg/ mL), the rate of Oxt infusion, and the frequency at which the rate was changed. An Oxt concentration of 50 mcg/mL gave the most consistent results with respect to the timing of birth, maternal care, and the survival of pups. This dosing paradigm was chosen after an iterative process involving adjustments in either the concentration of Oxt or the rate or frequency of administration. Approximately 7 days after delivery of the pups, the pump was removed from the subcutaneous pocket and the surgical site was restitched. These sutures were subsequently removed one week later after ensuring complete healing and closure. Higher concentration of Oxt (≥ 100 mcg/mL) or increased frequency of rate change was associated with dystocic labor and occasional cannibalization of the pups by the mother, though not to the same extent as pump implantation at E20.

**Western blotting for OxtR**

Approximately 10 μg of protein per lane was electrophoresed and transferred to membrane using Bolt western blot reagents (bolt 4-12% Bis Tris gel, catalog # NW04125; bolt sample reducing agent, catalog B0009; bolt LDS sample buffer, catalog # B0007; iBlot2 dry blotting system). The membrane was subsequently blocked with TBST buffer (catalog # S1012, EZ Bioresearch) containing 5% milk for 1 h at room temperature on a shaker. Following a brief wash with TBST buffer, the membrane was immunoblotted overnight at 4 °C on a shaker with primary rabbit anti-OxtR antibody (catalog # TA351476, OriGene Technologies), at a dilution of 1:250 in 5% milk-TBST buffer. HRP-conjugated secondary antibody (anti-rabbit IgG, catalog #7074, Cell Signaling Technology, Inc.) was used at a dilution of 1:1000 for 1 h at room temperature on a shaker. Immunoblots were incubated with ProSignal^®^ Dura ECL Reagent (catalog #20-301, Prometheus Protein Biology Products) for 2 minutes at room temperature and detection of bound antibody was achieved with LI-COR Fc imaging system (LI-COR Biosciences Inc.) and the band concentrations were analyzed with Image Studio Ver. 5.2. For loading control, the membrane was stripped using Western ReProbe PLUS (catalog# 786-307, G-Biosciences, Inc) according to manufacturer’s instructions and immunoblotted with HRP-conjugated β-Actin (mouse monoclonal IgG), at a dilution of 1:1000 (catalog # sc-47778, Santa Cruz Biotechnology Inc.) for 1 h and developed as described above.

**RNA-seq analysis of the newborn cortical transcriptome**

Samples were prepared according to library kit manufacturer’s protocol, indexed, pooled, and sequenced on an Illumina HiSeq. Basecalls and demultiplexing were performed with Illumina’s bcl2fastq software and a custom python demultiplexing program with a maximum of one mismatch in the indexing read. RNA-seq reads were then aligned to the Rattus norvegicus Ensembl Rnor_5.0 top-level assembly with STAR version 2.0.4b. Gene counts were derived from the number of uniquely aligned unambiguous reads by Subread: featureCount version 1.4.5. All gene counts were then imported into the R/Bioconductor package EdgeR and TMM normalization size factors were calculated to adjust for samples for differences in library size. Ribosomal genes and genes not expressed in at least 4 samples greater than one count-per-million were excluded from further analysis. The TMM size factors and the matrix of counts were then imported into the R/Bioconductor package Limma. Unknown latent effects were then estimated with the R/Bioconductor package SVA. Weighted likelihoods based on the observed mean-variance relationship of every gene and sample were then calculated for all samples with voomWithQualityWeights. Differential expression analysis was then performed to analyze for gene expression differences between conditions with Limma and perturbations in expression in known Gene Ontology (GO) terms and KEGG pathways with the R/Bioconductor package. The results were then filtered for only those genes or terms with Benjamini-Hochberg false-discovery rate adjusted p-values less than or equal to 0.05.

**2. Supplementary Figures**

**Fig. S1.** RNA-seq data showing the heatmap of differentially expressed genes after *in utero* exposure to anesthesia and surgery for pump implantation.

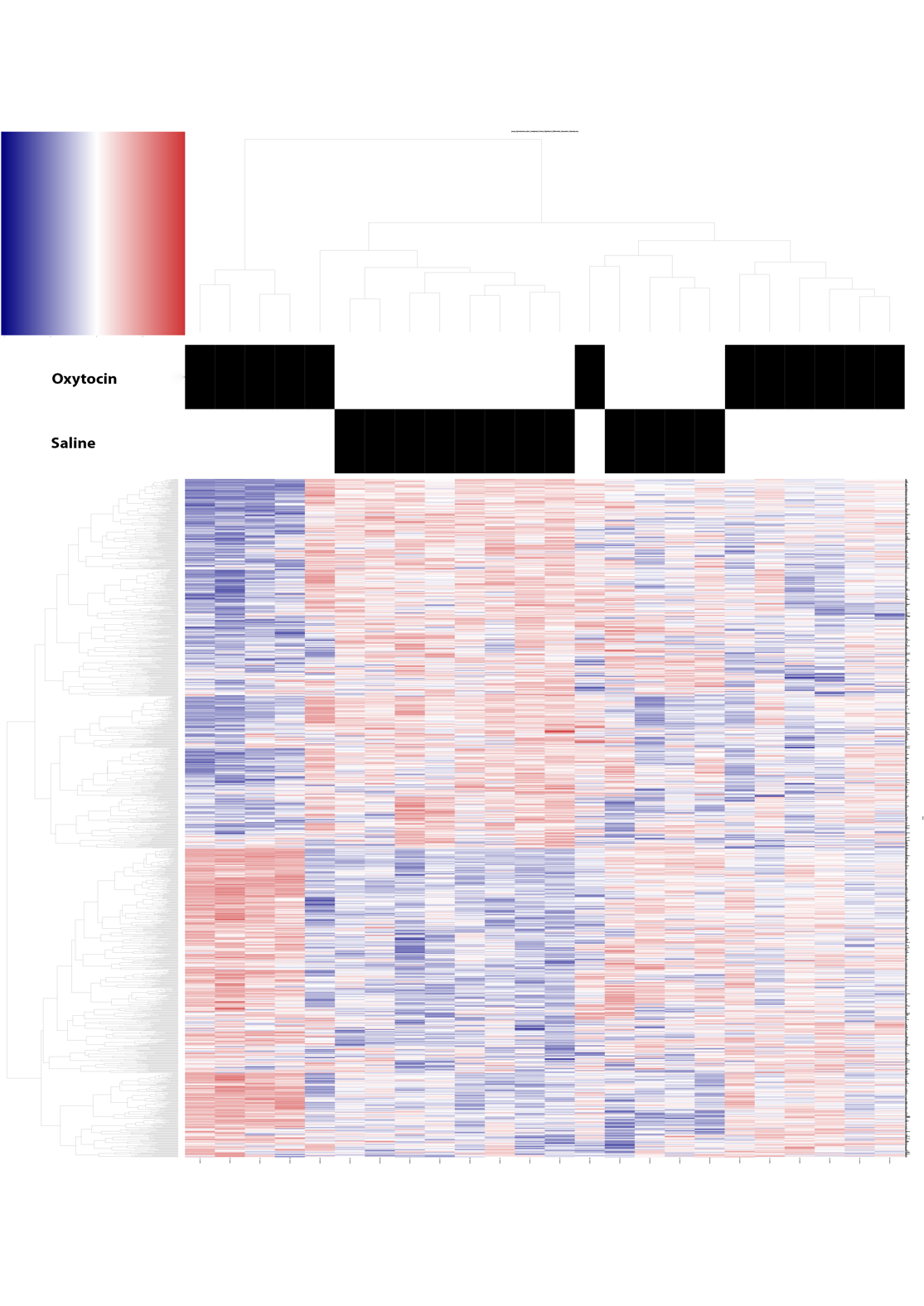

**Fig. S1. Differential gene expression in the developing brain after in utero exposure to anesthesia and surgery.** Overall, 12890 genes were differentially expressed of which 1180 genes were significantly differentially expressed with unadjusted p value < 0.05 (Supplementary Data S1 and S2). However, none of the genes were differentially expressed after adjustment for the false discovery rate.

**3. Supplementary Tables**

**Table S1.** **Animal use data**

| **Animal use** | **Age at pump implantation** | **Experiment** | **Outcome** |
| --- | --- | --- | --- |
| 4 | E 20 | Practice experiments to assess impact of anesthesia and surgery during pump implantation | High incidence of either preterm labor, dystocia, or cannibalization; pump implantation surgery advanced to earlier gestation at E18 |
| 8 | E18 | To determine optimal Oxt dosing regimen | Higher concentration of Oxt (≥ 100 mcg/mL) or increased frequency of infusion rate change were associated with dystocia and occasional cannibalization; Final Oxt concentration and infusion rate/rate change frequency chosen |
| 3 | E18 | Saline pump implantation to examine if exposure to surgery and anesthesia affects the neonatal cortical transcriptome at birth (RNA-seq) | Surgery and anesthesia at E18 were not associated with significant changes in the neonatal cortical transcriptome |
| 3 | Not applicable | Spontaneously laboring controls for the experiment above |  |
| 2 | E18 | Uterine telemetry experiment with Oxt initiated at E20 (i.e., one day earlier) to minimize the chances of spontaneous labor | Confirmation of the rise of intrauterine pressure after Oxt initiation |
| 12 | E18 | Oxt vs. saline pump implantation (n=6 each), followed by Oxt initiation at 72 h | Collection of cerebral cortices from vaginally delivered newborn pups (n=2 pups from each dam) to assess oxidative stress; the remaining offspring were weighed until weaning at P21 to determine the impact of Oxt on maternal nurturing and weight trajectory of the offspring |
| 12 | E18 | Oxt vs. saline pump implantation (n=7 and 5, respectively), followed by Oxt initiation early am of E21 to facilitate sample collection during the day | Collection of uterine myometrium for OxtR gene expression studies, OxtR immunofluorescence, and OxtR western blot |

**Table S1. Details of animal use for the creation and validation of the labor induction with oxytocin model.** A detailed description of the iterative process involved in the creation of the model that involved determination of the day of surgery, oxytocin concentration, rate of administration and the frequency of rate change. During the model development phase of the study, we used 12 dams in an iterative process to determine the optimal gestational age for surgery and a contextually relevant dosing paradigm for Oxt, with the ultimate goal of ensuring predictable delivery of viable and thriving pups and a nurturing mother. E: embryonic day; P: postnatal day.

**Table S2.** Litter data and weight gain trajectory of the offspring.

|  | Average litter size | Average weight gain (male pups)  (grams) | | | Average weight gain (female pups)  (grams) | | |
| --- | --- | --- | --- | --- | --- | --- | --- |
|  |  | P7 | P14 | P21 | P7 | P14 | P21 |
| Saline | 8.6 ± 1.4 | 12.0 ± 0.7 | 27.2 ± 0.7 | 41.9 ± 3.0 | 12.2 ± 1.3 | 26.6 ± 0.7 | 42.2 ± 3.8 |
| Oxt | 8.2 ± 1.2 | 14.8 ± 0.2 | 30.1 ± 2.1 | 48.9 ± 3.2 | 14.9 ± 0.5 | 29.1 ± 3.0 | 47.1 ± 4.0 |

**Table S2. Litter data and weight gain trajectory of the offspring.** Data were collected from the cohort of dams treated with either Oxt or saline *in utero* for the oxidative stress experiments. The litter size includes litter data from all 12 dams, but weight gain trajectories for the offspring from birth to weaning (P7, P14, P21) were available only for pups from 3 dams per treatment condition. Oxt-exposed pups were heavier than their saline-exposed counterparts at all time points. These observations need further investigation because the groups were not adjusted for either litter size or the time of birth (i.e., most Oxt-exposed pups were typically delivered at least 6-12 hours before saline-exposed pups).

**Table S3.** Taqman qPCR probe list from ThermoFisher Scientific

| Gene | Taqman primers Cat # | Reporter dye | Formulation |
| --- | --- | --- | --- |
| 18S rRNA | Hs99999901_s1 | FAM-MGB | 20x |
| gapdh | Rn01775763_g1 | FAM-MGB | 20x |
| pgk1 | Rn01474008_gH | FAM-MGB | 20x |
| actb | Rn00667869_m1 | FAM-MGB | 20x |
| mt-cyb | Rn03296746_s1 | FAM-MGB | 20x |
| mt-nd2 | Rn03296765_s1 | FAM-MGB | 20x |
| mt-nd5 | Rn03296799_s1 | FAM-MGB | 20x |
| mt-co1 | Rn03296721_s1 | FAM-MGB | 20x |
| mt-atp8 | Rn03296716_s1 | FAM-MGB | 20x |
| cat | Rn00560930_m1 | FAM-MGB | 20x |
| gpx1 | Rn00577994_g1 | FAM-MGB | 20x |
| gpx4 | Rn00822100_gH | FAM-MGB | 20x |
| gsr | Rn01482159_m1 | FAM-MGB | 20x |
| nox3 | Rn00585380_m1 | FAM-MGB | 20x |
| nox4 | Rn00585380_m1 | FAM-MGB | 20x |
| prdx1 | Rn00821587_g1 | FAM-MGB | 20x |
| sod1 | Rn00566938_m1 | FAM-MGB | 20x |
| sod2 | Rn00690588_g1 | FAM-MGB | 20x |
| txnip | Rn01533891_g1 | FAM-MGB | 20x |
| txnrd2 | Rn00574868_m1 | FAM-MGB | 20x |
| oxtR | Rn00563503_m1 | FAM-MGB | 20x |

4. Supplementary Movies

Movie S1. Experimental set up for iPRECIO^®^ pump implantation. A video walk-through of the overall surgical set up for performing the experiments.

Movie S2. Appropriate and nurturing care of the newborn after low dose (50 mcg/mL) Oxt regimen.

Movie S3. Poor maternal self-care and pup neglect in a dam implanted with iPRECIO^®^ pump at E20 and treated with high dose of Oxt (100 mcg/mL).

5. Supplementary Data Files

Data S1. RNA-seq data showing the list of significantly expressed genes in the developing cortex after *in utero* exposure to anesthesia and surgery.

Data S2. RNA-seq data showing the list of false discovery rate-unadjusted significantly expressed genes in the developing cortex after *in utero* exposure to anesthesia and surgery.
